# It is Never as Good the Second Time Around: Brain Areas Involved in Salience Processing Habituate During Repeated Drug Cue Exposure in Methamphetamine and Opioid Users

**DOI:** 10.1101/2020.04.18.036368

**Authors:** Hamed Ekhtiari, Rayus Kuplicki, Robin P Aupperle, Martin P. Paulus

**Author notes:** Corresponding Author: Hamed Ekhtiari, MD, PhD, Laureate Institute for Brain Research, 6655 South Yale Ave. Tulsa, OK 74136.

## Abstract

**Introduction:** The brain response to drug-related cues is an important marker in addiction-medicine, however, the temporal dynamics of this response in repeated exposure to the cues are not well known yet. In an fMRI drug cue-reactivity task, the presence of rapid habituation or sensitization was investigated by modeling time and its interaction with condition (drug>neutral) using an initial discovery-sample. Replication of this temporal response was tested in two other clinical populations.

**Methods:** Sixty-five male participants (35.8±8.4 years-old) with methamphetamine use disorder (MUD) were recruited as the discovery-sample. A linear mixed effects model was used to identify areas with a time-by-condition interaction in the discovery-sample. Replication of these effects was tested in two other samples (29 female with MUD and 22 male with opioid use disorder). The second replication-sample was re-tested within two weeks.

**Results:** In the discovery-sample, clusters within the VMPFC, amygdala and ventral striatum showed both significant condition and condition-by-time interaction with a habituation response for the drug-related cues but not neutral cues. The estimates for the main effects and interactions were generally consistent between the discovery and replication-samples across all clusters. The re-test data showed consistent lack of drug>neutral and habituation response within all selected clusters in the second cue-exposure session.

**Conclusions:** VMPFC, amygdala and ventral striatum show a habituation in response to drug-related cues which is consistent among different clinical populations. Habituation in response in the first session of cue-exposure and lack of reactivity in the second session of exposure provide foundations for development of cue-desensitization interventions.

## 1. Introduction

Drug craving has been considered as a motivational state that activates drug seeking behavior and potentiates relapse (H. Ekhtiari, Nasseri, Yavari, Mokri, & Monterosso, 2016; Wise, 1988) and is among one of the diagnostic criteria for substance use disorders in DSM 5 (American Psychiatric Association, 2013). Drug craving and subsequent relapse can be triggered by exposure to conditioned drug related cues whether they are pictures, mental images, smells, words, or actual drugs and their paraphernalia (Hamed Ekhtiari, Alam-Mehrjerdi, Nouri, George, & Mokri, 2010). In this context, drug cue exposure as an ecologically valid experimental paradigm has been developed to examine the mechanistic and predictive value of subjective, behavioral and biological responses to drug cues (Carter & Tiffany, 1999; Drummond, 2000). FMRI drug cue reactivity (FDCR) explores the neural response to drug cues presented to subjects inside the scanner (Hamed Ekhtiari, Faghiri, Oghabian, & Paulus, 2016). As of 2019, there are over 300 original published studies with FDCR. These studies reported positive findings that FDCR in certain brain areas is associated with severity of the drug addiction (Sjoerds, van den Brink, Beekman, Penninx, & Veltman, 2014; Smolka et al., 2006), defines prognosis of long-term recovery (Janes et al., 2010; Kosten et al., 2006), predicts the response to specific interventions (Courtney, Schacht, Hutchison, Roche, & Ray, 2016; Mann et al., 2014) and can be used as a proxy measure for effectiveness of different interventions in clinical trials (Lukas et al., 2013; Sadraee, Paulus, & Ekhtiari, 2019).

In fMRI drug cue reactivity studies, conventionally, the average response to drug cues in contrast to control conditions (neutral cues, fixation cross etc.) is reported/utilized as the main contrast/signal of interest (Hartwell et al., 2011; Schacht, Anton, & Myrick, 2013; Van Hedger, Keedy, Mayo, Heilig, & de Wit, 2018). However, drug cue reactivity is basically triggering a motivational/affective response that has a temporal behavior (H. Ekhtiari et al., 2016). With contemporary methodological conventions for fMRI cue reactivity tasks, there are multiple exposures to drug cues as blocks or events within each experiment. This provides a great opportunity to include time in the analysis models to explore this temporal behavior of response to drug cues. We expect to have a habituation response in some areas and also potentiation of response in some other areas during repeated trials of cue exposure. The dynamic response to affective cues have been reported previously in different brain areas including amygdala, ventral striatum and prefrontal cortices (Phan, Liberzon, Welsh, Britton, & Taylor, 2003; Wright et al., 2001). It has been reported frequently in fMRI studies that the amygdala habituates to the emotionally salient stimuli (Breiter et al., 1996; Fischer et al., 2003; Sladky et al., 2012) and its habituation is a more reliable index than its mean signal (Plichta et al., 2014). The ventral striatum also habituates in multiple exposures to rewards (Moses-Kolko et al., 2011). The first fMRI drug cue reactivity study that explored the temporal response to drug cues in the amygdala, ventral striatum, ventromedial prefrontal cortex (VMPFC) and anterior cingulate cortex, implemented prolonged exposure to drug cues (10 minutes videos) with fMRI and subjective measurements during and after the exposure (Murphy et al., 2017) and reported a correlation between the temporal response in the amygdala and subjective report of craving. Previous investigations had not considered the possibility of habituation/sensitization to the cues and therefore did not examine the effects of repeated presentation of drug cues on brain activation, however, we expect areas that are involved in reward and saliency processing present a time by condition (drug>neutral) interaction as their temporal dynamic response to the repeated presentation of drug cues.

This investigation examined three major questions. First, what is the time course of brain activation to repeated exposure of drug cues? Second, is there individual variability of the time course of brain activation that can be related to clinical characteristics, i.e., duration of dependence or abstinence? Third, what is the between-session reliability of the time course of brain activation in repeated drug cue exposure? To provide methodological resources to answer these questions, we have recently developed and validated a large pictorial database of drug and controlled neutral cues with 320 images and a series of fMRI drug cue reactivity tasks for opioid and methamphetamine users (Hamed Ekhtiari, Kuplicki, Pruthi, & Paulus, 2020). Having 8 blocks of drug or neutral cues makes it possible to explore the temporal dynamic response to the drug cues with adding time (blocks) as a factor in the analysis models. This experimental resource also provides an opportunity to test and compare drug cue reactivity in both opioid and methamphetamine users between or within different sample populations multiple times with distinct but equivalent drug cue sets. To answer questions one and two, we have recruited a discovery sample from methamphetamine users and conducted both whol-brain and ROI-based analysis to identify whether there are brain regions exhibiting time by condition (drug>neutral) interactions. Using clusters identified in this discovery sample, we examined whether the time by condition interaction effects replicated in two other replication samples, one with methamphetamine use disorder, but in a different clinical setting with longer duration of abstinence, and the other with opioid use disorder within the same clinical setting and same duration of abstinence as the discovery sample. To answer the third question, the second group was re-tested with a distinct but equivalent set of cues in the fMRI task within two weeks (Figure 1).

**Figure 1.**
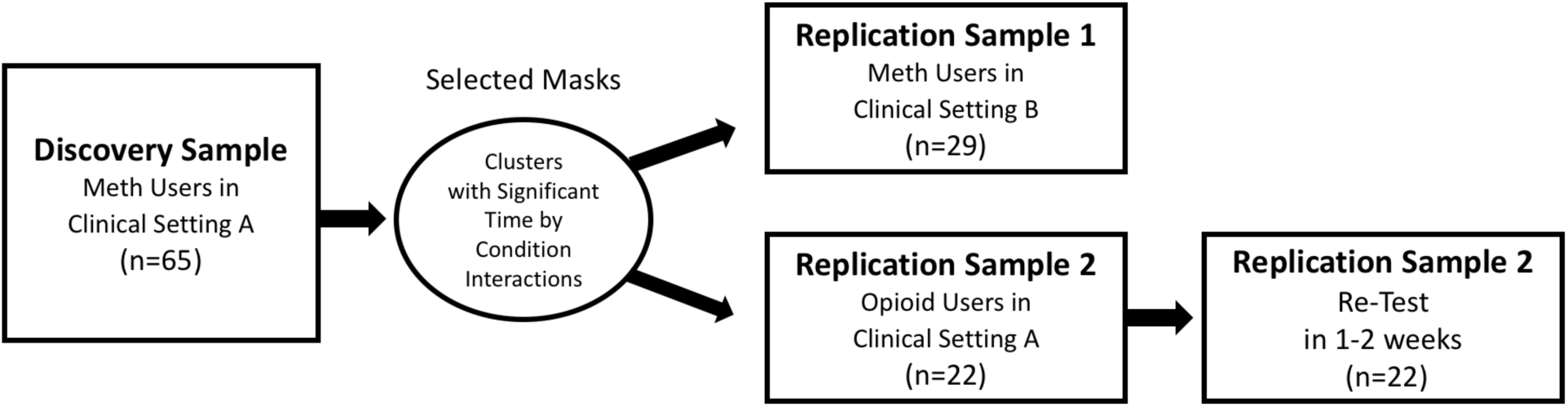
The study procedure with a discovery sample and two replication samples. Clusters with significant time by group interactions in both ROI and whole brain analysis in the discovery sample were tested in the replication samples.

## 2. Methods

In this study, the dynamic response to drug cues is explored in a discovery sample using an fMRI drug cue reactivity (FDCR) task and then the replication of the results was tested in two independent samples. The second replication sample was re-tested within two weeks with a distinct but equivalent version of the FDCR task to test the replication of the results within a sample population (Figure 1).

### 2.1. Participants

#### 2.1.1. Discovery Sample

The discovery sample (n=65) is recruited from men 35.86 years old (standard deviation (SD): 8.47) with methamphetamine use disorder who were admitted to a residential abstinence-based treatment center (12&12 center) in Tulsa, Oklahoma and were abstinent in the residential program or its aftercare transitional living programs. Inclusion criteria were (1) English speaking, (2) diagnosed with methamphetamine use disorder (last 12 months), (3) being abstinent from methamphetamine for at least one week and (4) willing and capable of interacting with the informed consent process. Exclusion criteria were (1) unwillingness or inability to complete any of the major aspects of the study protocol, including magnetic resonance imaging (i.e., due to claustrophobia), drug cue rating, or behavioral assessment, (2) abstinence from methamphetamine for more than 6 months based on self-report, (3) schizophrenia or bipolar disorder based on the MINI interview, (4) active suicidal ideation with intent or plan determined by self-report or assessment by PI or study staff during the initial screening or any other phase of the study, (5) positive drug test for amphetamines, opioids, cannabis, alcohol, phencyclidine, or cocaine confirmed by breath analyzer and urine tests.

#### 2.1.2. First Replication Sample

The first replication sample (n=29) was recruited from women with methamphetamine use disorder from the Women in Recovery (WIR) program at Family and Children’s Services located in Tulsa, Oklahoma program. WIR is an intensive outpatient “alternative to punishment” program for eligible women facing long prison sentences for non-violent drug-related offenses. The inclusion/exclusion criteria for recruitment were the same as the discovery sample, however the recruited sample is 1 year less educated (11.10 (SD:1.92) years compared to 12.12 (SD:1.81) years of education in the discovery sample). They have also been abstinent for a longer period time (112.73 (SD:49.21) days compared to 61.28 (SD: 36.29) days of abstinence in the discovery sample) (Table 1). In the MINI assessments, the first replication sample reported opioid use disorder and alcohol use disorder in a significantly lower level compared to the discovery sample (Table 1).

**Table 1.** Demographic and substance use profile of the three sample populations. Values are reported with mean (standard deviation) or frequency (percent%) format. Two replication samples were compared with the discovery sample with t-test and chi-square and the p values are reported in a column right of the replication sample values.

| Variables | Discovery Sample<br>(n=65) | Replication Sample 1<br>(n=29) | P Value | Replication Sample 2<br>(n=22) | P Value |
| --- | --- | --- | --- | --- | --- |
| Age (years) | 35.86 (8.47) | 32.44 (7.50) | 0.058 | 34.4 (6.26) | 0.410 |
| Education (years) | 12.13 (1.81) | 11.10 (1.92) | **0.010** | 12.90 (1.44%) | 0.059 |
| Substance Use Disorder Diagnoses (DSM V) |  |  |  |  |  |
| Stimulant (Amphetamines) Use Disorder | 65 (100%) | 29(100%) | 1.000 | 18 (81.82%) | 0.080 |
| Opioid Use Disorder | 28 (43.07%) | 2 (6.9%) | **0.001** | 22 (100%) | **<0.001** |
| Alcohol Use Disorder | 39 (60%) | 4 (13.79%) | **<0.001** | 6 (27.27%) | **0.016** |
| Cocaine Use Disorder | 6 (9.23%) | 0 (0%) | 0.966 | 2 (9.09%) | 1.000 |
| Cannabis Use Disorder | 30 (46.15%) | 19 (65.51%) | 0.131 | 10 (45.45%) | 1.000 |
| Sedative Use Disorder | 14 (21.54%) | 3 (10.34%) | 0.311 | 6 (27.27%) | 0.795 |
| Hallucinogen Use Disorder | 4 (6.15%) | 1 (3.45%) | 0.966 | 1 (4.55%) | 1.000 |
| Inhalant Use Disorder | 1 (1.54%) | 0(0%) | 1.000 | 0 (0%) | 1.000 |
| Duration of Current Abstinence (days) | 61.28 (36.29) | 112.73 (49.21) | **<0.001** | 50.25 (30.59) | 0.250 |

#### 2.1.3. Second Replication Sample

The second replication sample (n=22) is recruited from men with opioid use disorder who are admitted to the same clinical setting as the discovery sample (12&12 residential abstinence-based center in Tulsa, Oklahoma) with the same inclusion/exclusion criteria except for being diagnosed with opioid use disorder (last 12 months) without significant difference in age, education and duration of abstinence (Table 1). The second replication sample was recruited after the termination of recruitment for the discovery sample without any time overlap in the recruitment process.

### 2.2. Procedure

All three groups of recruited samples in this study were tested with the fMRI drug cue reactivity task introduced below at Laureate Institute for Brain Research (LIBR), Tulsa, OK. Data collection in these three groups was conducted as the baseline and pre-intervention assessments in three randomized clinical trials at LIBR between January 2018 and December 2019 (discovery sample from ClinicalTrials.gov Identifier: NCT03382379, replication sample 1 from NCT03922646 and replication sample 2 from NCT03907644). All participants signed written IRB approved consent forms and were assessed comprehensively including psychiatric assessments with DSM-V Mini-International Neuropsychiatric Interview (MINI) (Sheehan et al., 1998).

### 2.3. fMRI Task

All three groups of recruited samples in this study were tested with the same fMRI drug cue reactivity task structure (Hamed Ekhtiari et al., 2020). Pictorial cues within the FDCR task were selected from opioid or meth pictures based on the group of participants. In the fMRI task, participants are presented with blocks of cues that are either drug-related or neutral. Each block contains six images presented for five seconds each, with 200ms of blank screen between cues so that an entire block lasts 31 seconds. After every block, participants are asked to rate their current urge to use drug on a one to four scale, with one being “No Urge” and four being “Strong Urge”. The time between blocks varied between 8 and 12 seconds, and blocks alternated between drug and neutral, starting with neutral. A total of 8 blocks were presented, four of each condition. Total scan time for this task was approximately 6.5 minutes (Figure 2).

**Figure 2.**
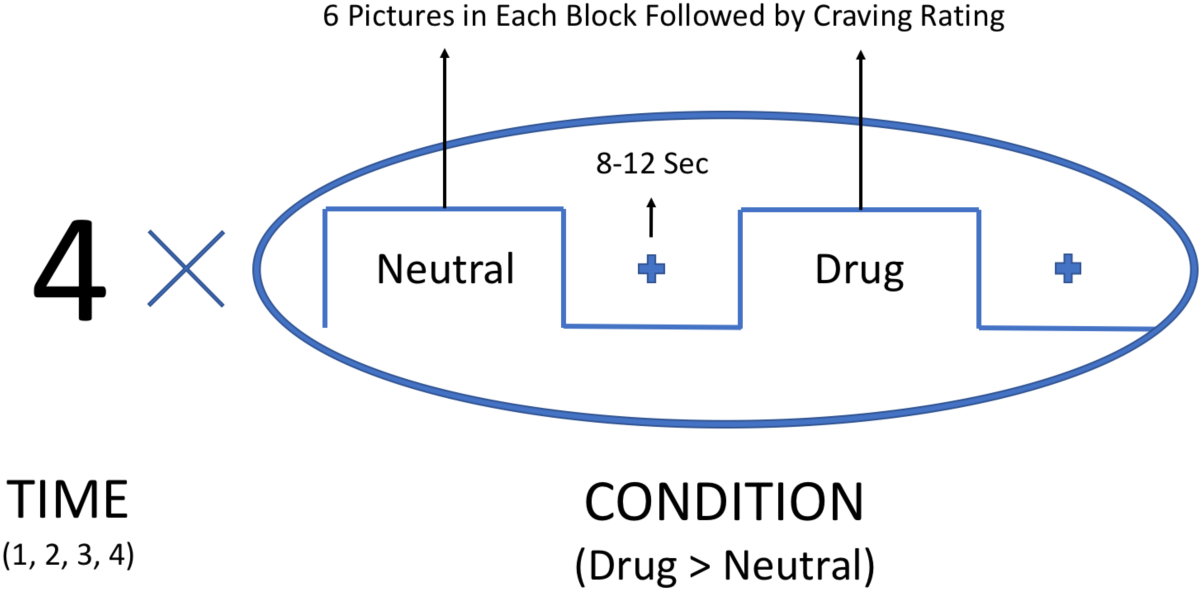
FMRI drug cue reactivity task structure and timings. There are 6 pictures within each block. Each picture is presented for 5 seconds with 0.2 seconds interstimulus interval. Each block is followed by a craving rating lasting 5 seconds. There is an inter-block interval of 8-12 seconds.

### 2.4. fMRI Data

MR images were acquired using two GE MRI 750 3T scanners at the LIBR. The FDCR task took 6 minutes 32 seconds of scan time with the following parameters: TR/TE=2000/27ms, FOV/slice=240/2.9mm, 128×128 matrix producing 1.875×1.875.2.9mm voxels, 39 axial slices, and 196 repetitions. High-resolution structural images were acquired through an axial T1-weighted magnetization-prepared rapid acquisition with gradient-echo (MPRAGE) sequence, using the following parameters: TR/TE=5/2.012ms, FOV/slice=240×192/0.9mm, 256×256 matrix producing 0.938×0.928×0.9mm voxels, and 186 axial slices.

### 2.5. Data Analysis

#### 2.5.1. fMRI Data Preprocessing

First level processing was done in AFNI and included removal of the first three pre-steady state images, despiking, slice timing correction, realignment, transformation to MNI space, and 4mm of Gaussian FWHM smoothing. Then, regression was carried out including nuisance regressors for the first three polynomial terms and the six motion parameters. TRs with excessive motion (defined as the Euclidean norm of derivative of the six motion parameters being greater than 0.3) were censored during regression.

#### 2.5.2. Temporal Dynamic fMRI Analysis

In order to evaluate the temporal dynamics of the BOLD response, we included separate regressors for each block of images, so that each participant had eight beta coefficients of interest computed (four neutral and four drug). At the group level, data were fit using a linear mixed effect in R. The model was “beta ∼ condition * time + motion” with a random effect for subject. Condition was either drug or neutral, with neutral in the intercept, and time was treated as a continuous variable with integer values of 1 through 4 (for the first through fourth blocks of each image type) which were then mean centered. This voxel-wise whole-brain analysis was used to identify clusters with a significant time by condition interaction (p < 0.001). Clusters in a priori regions of interest (ROIs; amygdala and ventral striatum (VStr), defined as Brainnetome ROIs 211+213 for left and 212+214 for right amygdala and ROIs 219, 220, 223, 224 for the VStr including the nucleus accumbens and ventral caudate) (Fan et al., 2016) were identified using a threshold of p < 0.05.

Individual-level dynamic responses (habituation slopes) were estimated for each ROI exhibiting a condition by time interaction as well as self-reported craving. Slopes were computed by fitting separate linear models for each subject with beta ∼ condition * time + motion for BOLD responses and craving ∼ condition * time for self-reported craving. From each linear model, we took the beta for time in each condition (drug or neutral) to be the subject’s temporal response to that condition. The condition by time interaction is taken to be the subject’s selective habituation to drug related images, with negative values indicate a decreasing response to drug related images when compared to the change in response to neutral images.

#### 2.5.5. Replication Testing

The temporal response to four drug and neutral blocks within the selected masks (significant clusters) discovered from the time by condition (drug/neutral) interaction in the discovery sample was explored in two replication samples. We fit four independent LME (linear mixed effect) models in three sample populations (one assessed twice) for the average beta estimates within the masks. For model comparison, the estimates for the effect of condition and the condition by time interaction between each replication sample and the discovery sample were compared using Z-tests.

## 3. Results

### 3.1. Discovery sample results

#### 3.1.1. Main effect of condition

Using a whole brain analysis in the discovery sample, the main effect of cue type (condition: drug>neutral) is significant in areas reported in previous cue reactivity studies including the prefrontal cortex, amygdala, striatum, insula and secondary visual processing areas. The main effect of condition is depicted in figure 3 panel a.

**Figure 3.**
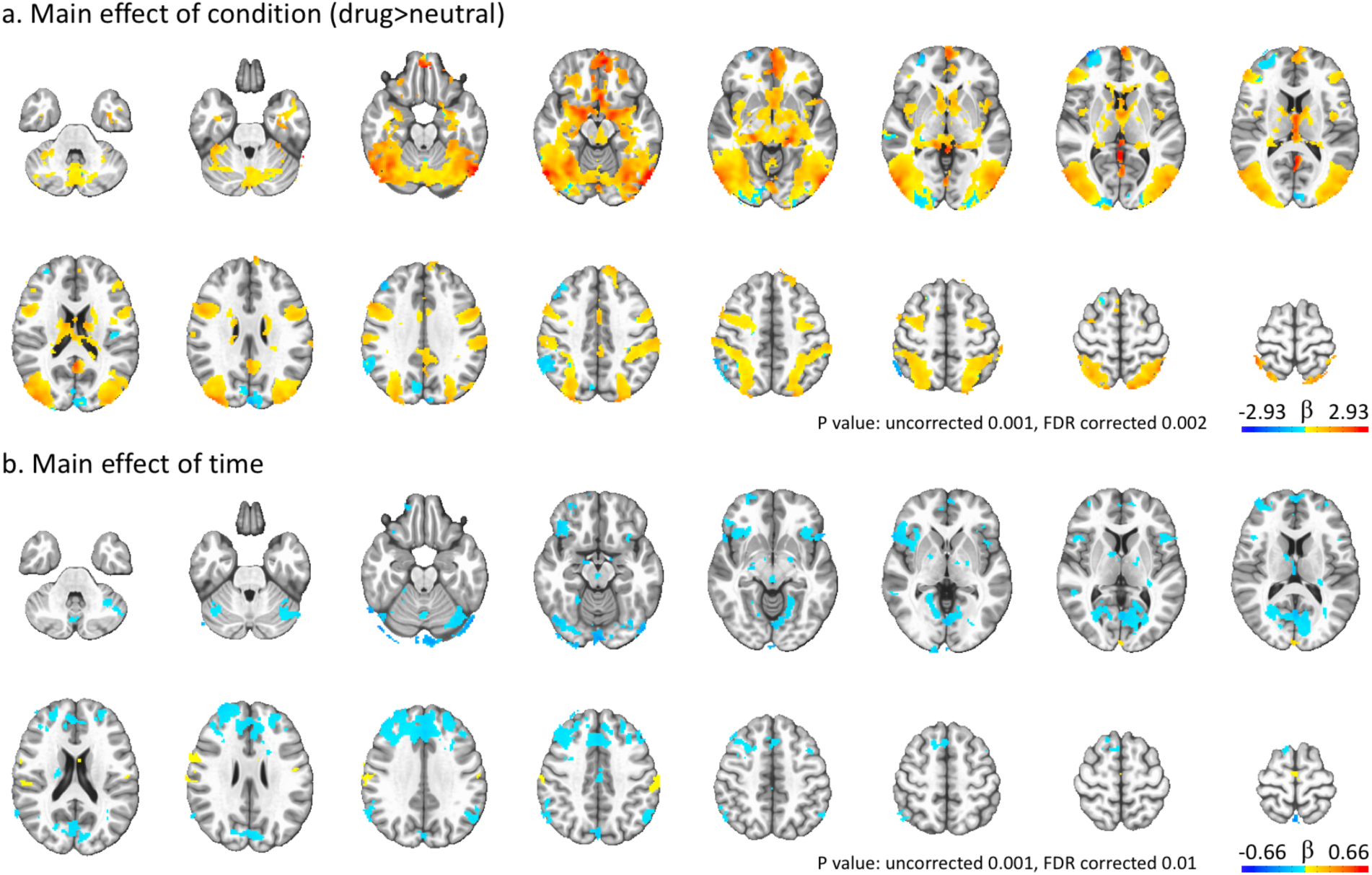
Whole brain response to the fMRI drug cue reactivity task in the discovery sample (n=65). Panel a. main effect of condition (craving>neutral) and b. main effect of time.

#### 3.1.2. Main effect of time

Several clusters in areas like the dorsolateral prefrontal (DLPFC), anterior insula cortex (AIC) and dorsal anterior cingulate cortex (dACC) show a negative main effect of time but no condition by time interaction, indicating attenuated responses to both categories of cues throughout the task. The main effect of time is depicted in figure 3 panel b.

#### 3.1.3. Condition*time interaction

Clusters within the ventromedial prefrontal cortex (VMPFC), bilateral superior temporal gyrus (STG) (Figure 4), right amygdala and bilateral ventral striatum (Figure 5) show a significant condition by time interaction with an attenuating response for the drug related cues (habituation). No area shows a significant condition by time interaction with an escalating response for the drug related cues (sensitization). Masks from these six clusters were generated to test the response within the replication samples.

**Figure 4.**
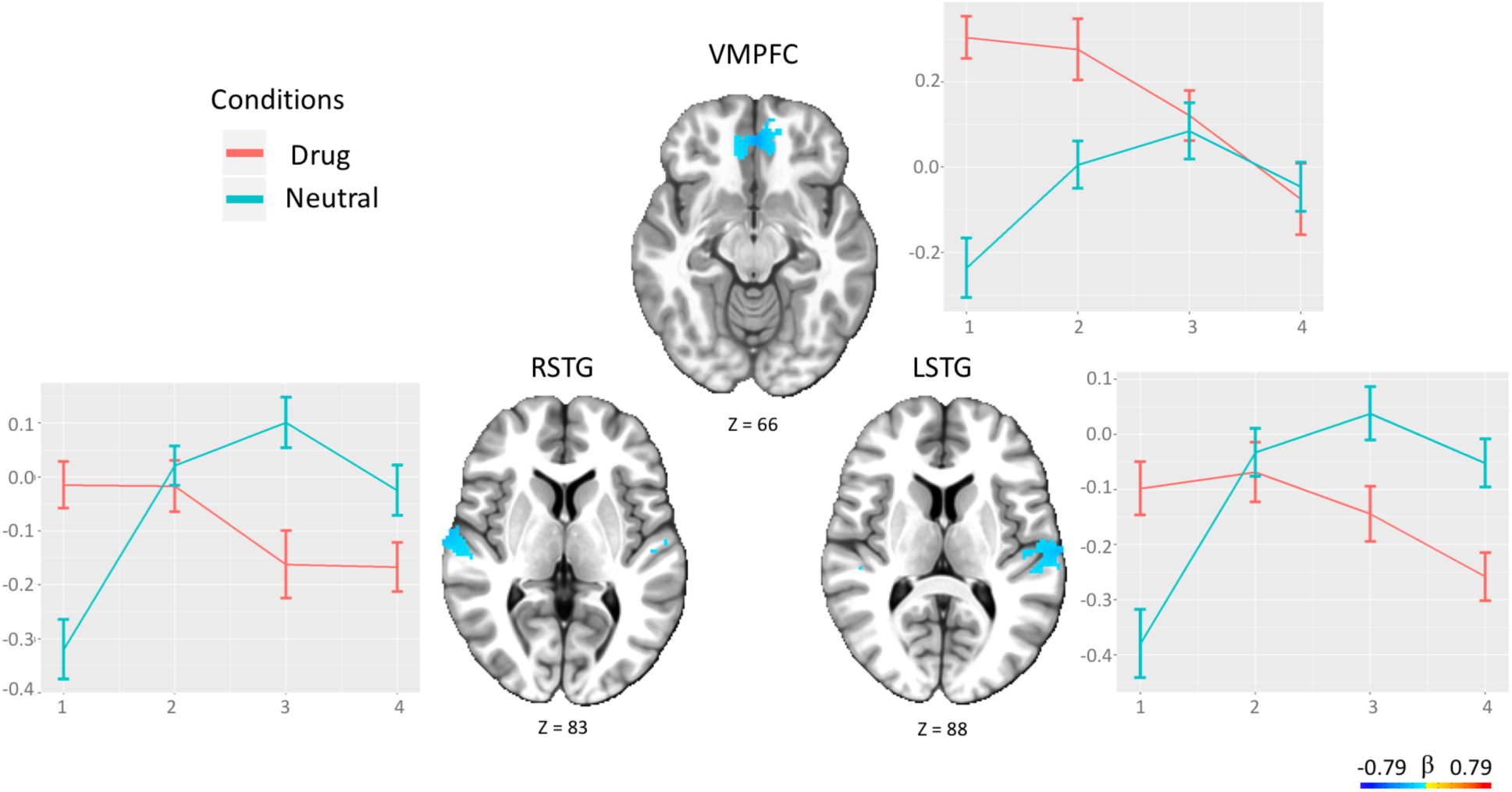
Temporal behavior of clusters with time by condition interaction in whole brain analysis in the discovery sample. VMPFC: ventromedial prefrontal cortex, RSTG: right superior temporal gyrus, LSTG: left superior temporal gyrus (p < 0.001).

**Figure 5.**
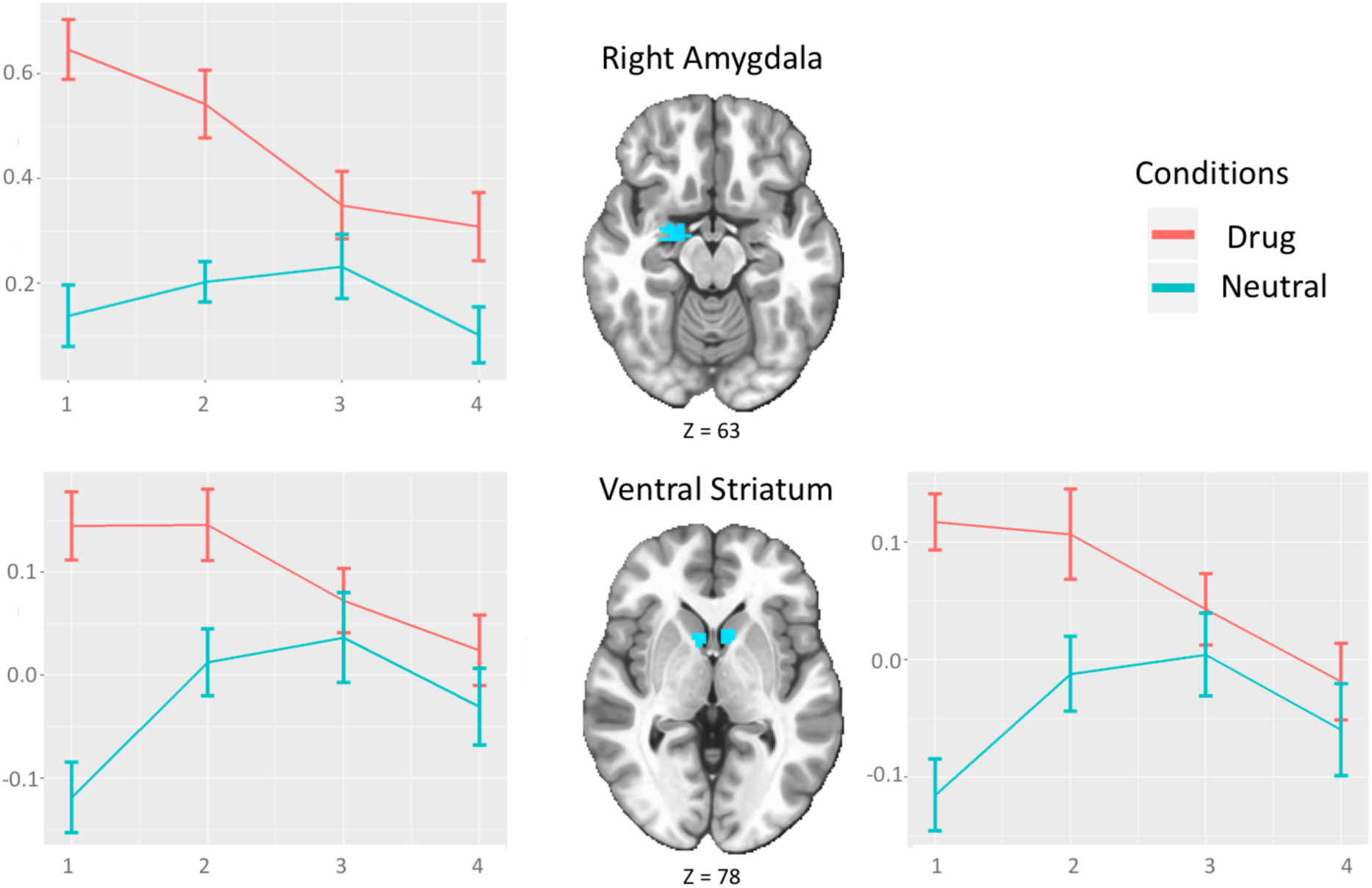
Temporal behavior of clusters with time by condition interaction within a priori ROIs (amygdala and ventral striatum, i.e., head of caudate and nucleus accumbens) (p <0.05)

#### 3.1.4. Individual habituation slopes

The correlation matrix between the clinical characteristics, i.e., duration of methamphetamine use, monthly cost of methamphetamine use, and duration of abstinence and craving self-reports before and after scanning (visual analogue scale 0-100) with individual habituation slopes within the six cluster masks and also subjective reports of craving inside the scanner after each block shows no significant correlation between the habituation slopes and clinical characteristics or craving self-reports outside the scanner that can survive p-value<0.05 threshold corrected for false discovery rate (FDR) (Figure S1).

### 3.2. Region of Interest Results

Independent LME models were tested in the discovery sample population and two replication samples (one assessed twice) for the average beta estimates within the six cluster masks generated from the discovery sample.

#### 3.2.1. VMPFC

In the discovery sample, the LME model within VMPFC mask shows highly significant estimate for the condition by time interaction (t=-4.97, p-value<0.001). There is also a significant effect of condition (t=4.55, p-value<0.001) and marginally significant effect of time (t=1.98, p-value=0.04). The condition by time interaction is significant in both replication sample 1 (t=-2.54, p-value= 0.01) and the first assessment of replication sample 2 (t=-3.24, p-value=0.01), but not in the retest in replication sample 2 (t=-0.35, p-value= 0.72) (Table 2 and Figure 6). The effect of motion is not significant in any of the 4 models. In the model comparison by contrasting the estimates for the condition by time interaction between each replication sample and the discovery sample using Z-tests, there is no significant difference (p-values > 0.45) between the discovery sample and replication samples 1 and 2; however there is a significant difference between the discovery sample and replication sample 2-retest (Z=2.52, p-value=0.033).

**Table 2.** Replication of Temporal Dynamic Response to Drug-related Cues Compared to Neutral Cues. Columns represent beta coefficients, standard errors (SEs) and p-values from four independent LME (linear mixed effect) models in three sample populations (one assessed twice) for the average beta estimates within the six masks obtained from the initial discovery phase. VMPFC: ventromedial prefrontal cortex, RSTG: right superior temporal gyrus, L VStr: left ventral striatum.

|  | VMPFC |  |  | RSTG |  |  | LSTG |  |  | R Amygdala |  |  | R VStr |  |  | LVStr |  |  |
| --- | --- | --- | --- | --- | --- | --- | --- | --- | --- | --- | --- | --- | --- | --- | --- | --- | --- | --- |
| Discovery Sample | Beta | SE | P value | Beta | SE | P value | Beta | SE | P value | Beta | SE | P value | Beta | SE | P value | Beta | SE | P value |
| Intercept | 0.057 | 0.071 | 0.425 | 0.006 | 0.051 | 0.906 | -0.008 | 0.05 | 0.868 | 0.223 | 0.069 | 0.002 | 0.056 | 0.034 | 0.109 | 0.095 | 0.036 | 0.01 |
| Motion | -0.992 | 0.582 | 0.093 | -0.53 | 0.42 | 0.211 | -0.801 | 0.407 | 0.054 | -0.568 | 0.576 | 0.327 | -0.995 | 0.28 | **0.001** | -1.216 | 0.292 | **<0.001** |
| Condition (drug>neutral) | 0.192 | 0.042 | **<0.001** | -0.037 | 0.031 | 0.228 | -0.044 | 0.032 | 0.169 | 0.292 | 0.035 | **<0.001** | 0.106 | 0.021 | **<0.001** | 0.12 | 0.023 | **<0.001** |
| Time | 0.051 | 0.026 | **0.048** | 0.095 | 0.02 | **<0.001** | 0.1 | 0.02 | **<0.001** | -0.005 | 0.022 | 0.818 | 0.017 | 0.014 | 0.202 | 0.027 | 0.015 | 0.062 |
| Condition:Time | -0.182 | 0.037 | **<0.001** | -0.153 | 0.028 | **<0.001** | -0.152 | 0.029 | **<0.001** | -0.105 | 0.031 | **0.001** | -0.064 | 0.019 | **0.001** | -0.072 | 0.021 | **0.001** |
| **Replication Sample 1** |  |  |  |  |  |  |  |  |  |  |  |  |  |  |  |  |  |  |
| Intercept | -0.049 | 0.088 | 0.579 | -0.139 | 0.095 | 0.156 | -0.17 | 0.104 | 0.113 | 0.166 | 0.089 | 0.07 | -0.14 | 0.062 | 0.031 | -0.14 | 0.083 | 0.104 |
| Motion | -0.025 | 0.751 | 0.974 | 1.175 | 0.832 | 0.169 | 1.862 | 0.899 | 0.048 | 0.022 | 0.765 | 0.978 | 0.612 | 0.543 | 0.27 | 0.953 | 0.734 | 0.205 |
| Condition (drug>neutral) | 0.187 | 0.06 | 0.004 | -0.172 | 0.052 | 0.003 | -0.182 | 0.064 | 0.005 | 0.246 | 0.055 | **<0.001** | 0.085 | 0.034 | 0.014 | 0.085 | 0.038 | 0.028 |
| Time | 0.084 | 0.038 | **0.027** | 0.055 | 0.03 | 0.069 | 0.079 | 0.04 | 0.052 | 0.048 | 0.029 | 0.099 | 0.016 | 0.022 | 0.446 | 0.053 | 0.024 | **0.031** |
| Condition:Time | -0.136 | 0.053 | **0.012** | -0.081 | 0.043 | 0.059 | -0.108 | 0.057 | 0.06 | -0.073 | 0.041 | 0.074 | -0.035 | 0.031 | 0.249 | -0.079 | 0.034 | **0.022** |
| **Replication Sample 2** |  |  |  |  |  |  |  |  |  |  |  |  |  |  |  |  |  |  |
| Intercept | 0.069 | 0.108 | 0.525 | 0.069 | 0.078 | 0.389 | 0.007 | 0.096 | 0.941 | 0.278 | 0.092 | 0.006 | 0.032 | 0.072 | 0.656 | 0.058 | 0.092 | 0.538 |
| Motion | -1.405 | 0.828 | 0.105 | -0.696 | 0.612 | 0.268 | -0.286 | 0.747 | 0.706 | -1.17 | 0.726 | 0.123 | -0.567 | 0.566 | 0.328 | -0.723 | 0.728 | 0.332 |
| Condition (drug>neutral) | 0.145 | 0.076 | 0.058 | -0.06 | 0.048 | 0.218 | -0.113 | 0.06 | 0.063 | 0.145 | 0.049 | **0.004** | 0.072 | 0.041 | 0.08 | 0.108 | 0.051 | **0.036** |
| Time | 0.066 | 0.048 | 0.174 | 0.091 | 0.031 | **0.003** | 0.131 | 0.038 | **0.001** | 0.018 | 0.031 | 0.564 | 0.008 | 0.026 | 0.75 | 0.032 | 0.032 | 0.331 |
| Condition:Time | -0.22 | 0.068 | **0.001** | -0.117 | 0.043 | **0.007** | -0.176 | 0.054 | **0.001** | -0.126 | 0.044 | **0.004** | -0.096 | 0.037 | **0.009** | -0.121 | 0.046 | **0.009** |
| **Replication Sample 2-Retest** |  |  |  |  |  |  |  |  |  |  |  |  |  |  |  |  |  |  |
| Intercept | 0.078 | 0.127 | 0.542 | -0.031 | 0.135 | 0.823 | -0.036 | 0.148 | 0.81 | 0.249 | 0.117 | 0.043 | -0.151 | 0.083 | 0.083 | -0.063 | 0.084 | 0.46 |
| Motion | -0.274 | 1.225 | 0.825 | 0.689 | 1.322 | 0.608 | 0.315 | 1.47 | 0.832 | -0.102 | 1.121 | 0.928 | 1.202 | 0.813 | 0.155 | 0.54 | 0.824 | 0.52 |
| Condition (drug>neutral) | -0.019 | 0.072 | 0.789 | -0.152 | 0.064 | **0.028** | -0.124 | 0.057 | **0.03** | 0.076 | 0.071 | 0.295 | 0.02 | 0.039 | 0.61 | -0.006 | 0.04 | 0.875 |
| Time | 0.006 | 0.046 | 0.895 | -0.008 | 0.034 | 0.825 | 0.015 | 0.036 | 0.682 | -0.045 | 0.033 | 0.176 | -0.026 | 0.024 | 0.297 | -0.012 | 0.025 | 0.646 |
| Condition:Time | -0.023 | 0.065 | 0.721 | -0.017 | 0.048 | 0.72 | -0.022 | 0.051 | 0.66 | -0.023 | 0.047 | 0.618 | 0.028 | 0.035 | 0.428 | 0.027 | 0.036 | 0.451 |

**Figure 6.**
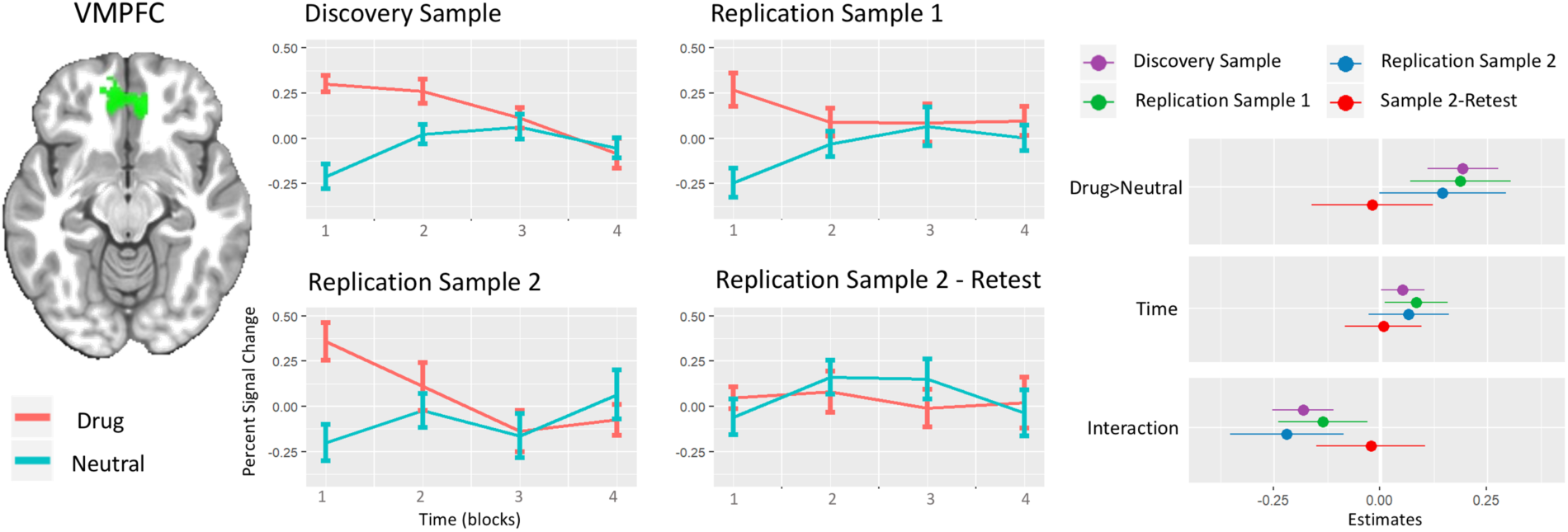
Replication of Temporal Dynamic Response to Drug-related Cues Compared to Neutral Cues within VMPFC. Middle panels show the temporal response to four drug and neutral blocks within the VMPFC mask discovered from whole brain time by condition (drug/neutral) interaction in the discovery sample; error bars represent +-one standard error. The right panels show coefficients from four independent LME (linear mixed effect) models in three sample populations (one assessed twice) for the average beta estimates within the VMPFC mask. Whiskers depict 95 percent confidence intervals. In the model comparison by contrasting the estimates for the effect of condition and the condition by time interaction between each replication sample and the discovery sample using Z-tests, there is no significant difference (p-values > 0.45) between the discovery sample and replication samples 1 and 2; however there is a significant difference between the discovery sample and replication sample 2-retest in both condition (drug>neutral)(p-value=0.011) and condition by time interaction (p-value=0.033).

#### 3.2.2. STG

There is no significant effect of condition in the STG in the discovery or replication samples 1 and 2. However, in the replication sample 2 re-test, there is a significant negative (neutral>drug) effect of condition in both masks (right: t=-2.35, p-value=0.028 and left: t=-2.18, p-value=0.03) (Figure S2). The LME model within both right and left STG masks shows a highly significant estimate for the condition by time interaction (right: t=-5.53, p-value<0.001 and left: t=-5.33, p-value<0.001). The condition by time interaction is significant in both masks in the replication sample 2 (right: t=-2.71, p-value=0.007 and left: t=-3.27, p-value=0.001) but not in the re-test in the same sample. In the replication sample 1 the condition by time interaction is marginally significant in both masks (right: t=-1.90, p-value=0.059 and left: t=-1.88, p-value=0.06) (Table 2 and Figure S2). There is no effect of motion in the STG masks. In the model comparison by contrasting the estimates for the condition by time interaction between each replication sample and the discovery sample using Z-tests, there is no significant difference (p-values>0.16) between the discovery sample and replication samples 1 and 2 in both masks; however the difference between the discovery sample and replication sample 2-retest is significant in both masks (right: Z=-2.44, p-value=0.01 and left: Z=-2.22, p-value=0.02).

#### 3.2.3. Amygdala

The discovery sample LME model within the amygdala mask shows a highly significant estimate for the condition by time interaction (t=-3.43, p-value=0.001). There is also a significant effect of condition (t=8.45, p-value<0.001). In three other replication samples, the effect of condition by time interaction is significant in replication sample 2 (t=-2.88, p-value= 0.004) and marginally significant in replication sample 1 (t=-1.80, p-value=0.07), but not in the retest of replication sample 2 (t=-0.5, p-value= 0.618) (Table 2 and Figure 7). The effect of motion is not significant in any of the 4 models for the amygdala mask. In the model comparison by contrasting the estimates for the condition by time interaction between each replication sample and the discovery sample using Z-tests, there is no significant difference (p-values > 0.14) between the discovery sample and replication samples 1 and 2 and replication sample 2-retest.

**Figure 7.**
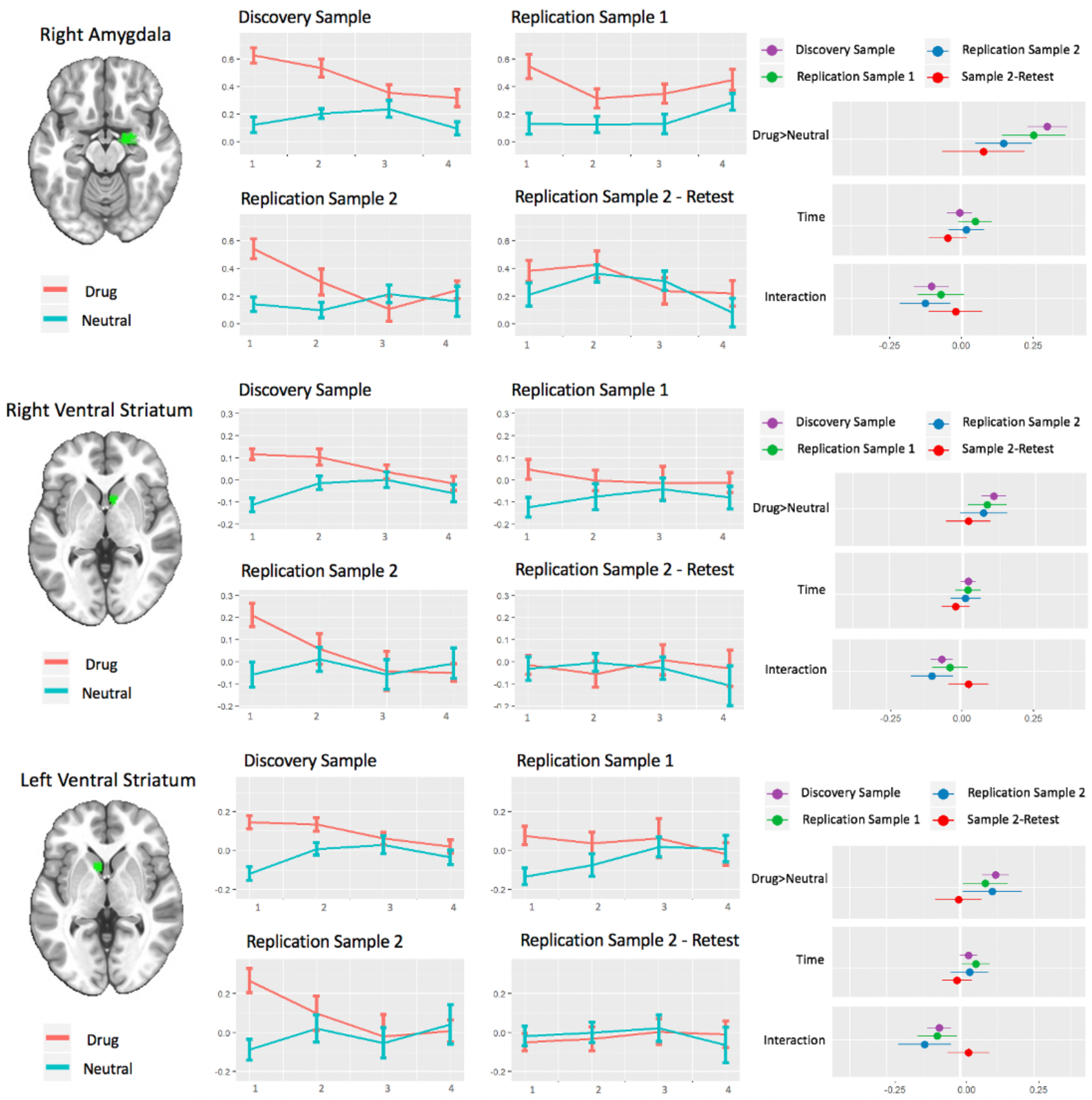
Replication of Temporal Dynamic Response to Drug-related Cues Compared to Neutral Cues within Amygdala and Ventral Striatum Masks. Middle panels show the temporal response to four drug and neutral blocks within the masks discovered from time by condition (drug/neutral) interaction in the discovery sample; error bars represent +-one standard error. The right panels show coefficients from four independent LME (linear mixed effect) models in three sample populations (one assessed twice) for the average beta estimates within the masks. Whiskers depict 95 percent confidence intervals.

#### 3.2.4. Ventral Striatum

The discovery sample LME model within both right and left VStr masks shows a highly significant estimate for the condition by time interaction (right: t=-3.31, p-value=0.001 and left: t=-3.47, p-value=0.001). There is also a significant effect of condition in both masks (right: t=4.95, p-value<0.001 and left: t=5.17, p-value<0.001). The condition by time interaction is significant in both masks in replication sample 2 (right: t=-2.63, p-value=0.009 and left: t=-2.64, p-value=0.009) but not in the re-test in the same sample. In the replication sample 1 the condition by time interaction is significant for the left mask (t=-2.3, p-value=0.02), but not in the right mask (Table 2 and Figure 7). The effect of motion is significant for both right and left masks just in the discovery sample (p-value=<0.001). In the model comparison by contrasting the estimates for the condition by time interaction between each replication sample and the discovery sample using Z-tests, there is no significant difference (p-values>0.32) between the discovery sample and replication samples 1 and 2 in both masks; however the difference between the discovery sample and replication sample 2-retest is significant in both masks (right: Z=-2.3, p-value=0.02 and left: Z=-2.54, p-value=0.01).

### 3.3. Subjective self-report of craving

There is a significant higher level of momentary craving (urge) self-report after seeing drug related blocks compared to the neutral blocks in all four groups (p-value<0.001 in all groups) (Figure 8). The difference in response to the drug related blocks compared to the neutral blocks were higher in the replication sample 2 and its retest (opioid cues) compared to two other groups (methamphetamine cues) (p-value<0.01 in all two groups comparisons). There is a small time by condition interaction in the discovery sample (Estimate=-0.1, p-value=0.014) that was not replicated in other groups. The individual habituation slope for the time by condition interaction in the discovery sample was not correlated with individual habituation slopes in the 6 cluster masks (Figure S1). There is no significant difference in response to the drug related blocks compared to the neutral blocks test in test-retest in the replication sample 2 (Z=-0.13, p-value=0.89) with a reasonable test-retest reliability (ICC=0.69, 95%CI 0.40-0.86).

**Figure 8.**
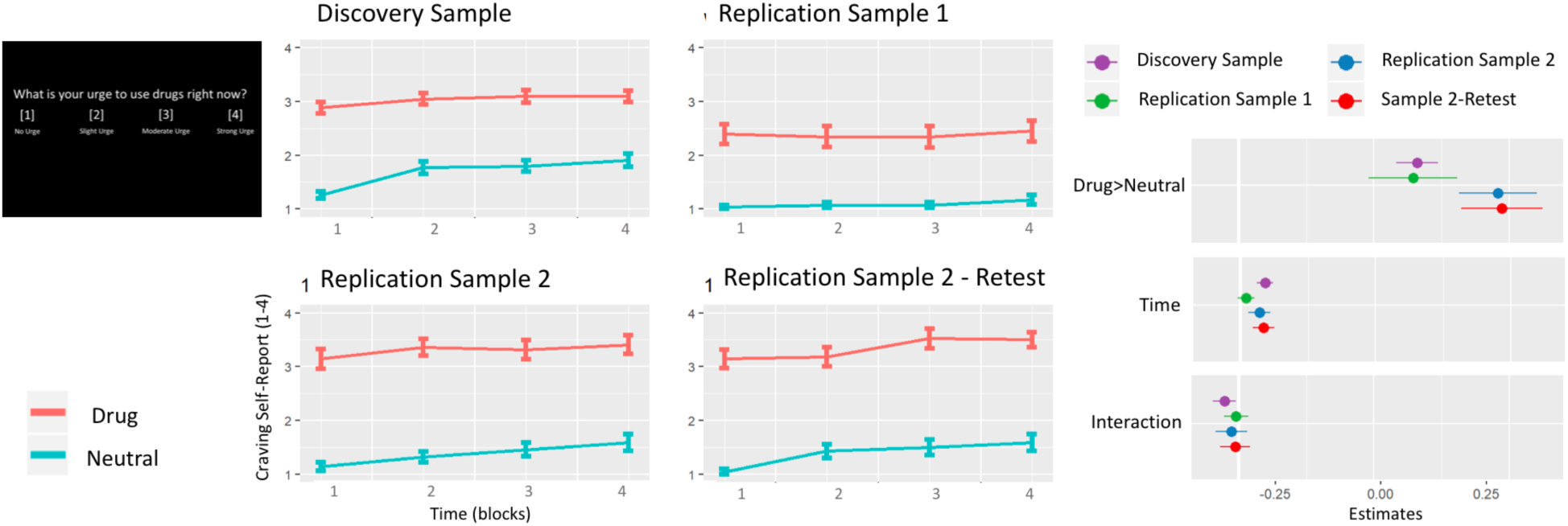
Subjective Report of Craving after Drug-related and Neutral Blocks. Middle panels show the subjective report of momentary craving (urge) after to four drug and neutral blocks in four levels, 1: no urge, 2: slight urge, 3: moderate urge and 4: strong urge; error bars represent +-one standard error. The right panels show coefficients from four independent LME (linear mixed effect) models in three sample populations (one assessed twice) for the subjective reports. Whiskers depict 95 percent confidence intervals.

## 4. Discussion

This investigation examining the time course of brain activation in response to drug cue exposure in a group of individuals with methamphetamine and opioid use disorder yielded three main results. First, we found VMPFC, right amygdala and bilateral ventral striatum respond to drug cues with higher activation compared to the neutral cues and then show a significant habitation in this response in the repeated cue presentation. We found that none of the brain areas shows an increased response to drug cues over time (sensitization). Second, we found the habituation response in these areas replicated consistently in independent samples of opioid and methamphetamine users. Third, we found that the areas exhibiting the habituation response in baseline sessions are not responsive to the drug cues in the second session of cue exposure few days later.

Habituation in response to repeated presentation of conditioned cues is a well-known phenomenon in the behavioral neuroscience in both animal and human models (Phelps, Delgado, Nearing, & LeDoux, 2004; Quirk & Mueller, 2008). The role of VMPFC, amygdala and ventral striatum, as the central hubs for dynamic processing the affective/reinforcing value of the environmental cues (Phan et al., 2003; Phan, Wager, Taylor, & Liberzon, 2002; Sescousse, Caldú, Segura, & Dreher, 2013), is reported frequently in the habituation response to the salient cues (Breiter et al., 1996; Fischer et al., 2003; Ishai, Pessoa, Bikle, & Ungerleider, 2004; Sladky et al., 2012; Strauss et al., 2005; Wright et al., 2001). Novelty detection, rapid processing of the saliency prediction errors and dynamic evaluation of the environment instead of a sustained processing can be foundation of this rapidly habituating response. Consistent with our findings, the right amygdala is shown to dynamically respond to the salient cues, while the left amygdala is specialized for sustained evaluations of the salient cues (Fischer et al., 2003; Wright et al., 2001)

The role of bilateral superior temporal gyrus (STG) in drug cue reactivity was reported in previous studies (Kang et al., 2012; McClernon, Kozink, & Rose, 2008; Yalachkov, Kaiser, & Naumer, 2012). In our study, STG does not show any positive or negative main effect of condition (drug>neutral), however, it consistently shows a significant time by condition interaction with a rapid attenuation in response to drug related cues compared to the neutral cues. The role of STG in visuotemporal attentional processing of environmental stimuli might be one of the explanations for this finding (Li, Chen, Han, Chui, & Wu, 2012). It is reported that lesion in STG is associated with more prolonged deployment of visuotemporal attention (Shapiro, Hillstrom, & Husain, 2002). Drug cues might engage STG as a rapidly responding area to attentionally salient stimuli, however, this attentional saliency will drop rapidly for drug cues and overall there is not a higher signal for drug cues compared to the neutral cues in the STG.

Habituation in response to repeated presentation of conditioned cues is a foundation for cue exposure therapy in psychiatric disorders like phobias and PTSD. Habituation in amygdala response to emotional cues is reported as an outcome of cue exposure therapy in people with phobia (Goossens, Sunaert, Peeters, Griez, & Schruers, 2007) and VMPFC activity during early extinction could predict the outcome of the cue exposure therapy in people with phobia (Lange et al., 2020). Furthermore, stimulating the VMPFC with repetitive transcranial magnetic stimulation (rTMS) in a placebo-controlled trial improved treatment outcome of the cue exposure therapy. Habituation in response to affective stimuli in VMPFC, amygdala and ventral striatum is also being targeted in the fMRI neurofeedback studies for reduction of affective response among people with depression (Young et al., 2014), PTSD (Nicholson et al., 2017; Zotev et al., 2018) and healthy participants (Haller et al., 2013; Paret et al., 2016; Zotev et al., 2011; Zotev, Phillips, Young, Drevets, & Bodurka, 2013).

Our study is one of the first fMRI studies reporting a habituation in response in the context of repeated drug cue exposure. Drug cue exposure therapies have not been as successful in addiction medicine as the cue exposure therapies in areas like phobias or PTSD (Marissen, Franken, Blanken, van den Brink, & Hendriks, 2007; Martin, LaRowe, & Malcolm, 2010; Mellentin et al., 2017). The only published study on the brain response to a cue exposure therapy intervention among drug user, reported higher reduction in response to alcohol cues in few clusters across the brain including ventral striatum after nine sessions of cue exposure in people with alcohol use disorder (Vollstädt-Klein et al., 2011). We found in this study that from many areas of activation due to the exposure to the drug cues, only VMPFC, amygdala and ventral striatum showed any habituation in response to repeated exposure to the drug cues in the temporal window of our fMRI task. It can be expected that conventional cue exposure interventions will cause habituation and learning extinction across few neuro/cognitive processes from many that are involved in drug craving (Byrne, Haber, Baillie, Giannopolous, & Morley, 2019; H. Ekhtiari et al., 2016).

We have not found any consistent habituation or sensitization in the subjective craving responses after being exposed to the drug blocks compared to the neutral blocks. We have also not found any significant correlation between craving self-reports and habituation slopes in the six cluster masks. Phan et. al., 2003 have also reported lack of time by condition interaction in the subjective reports to the emotionally salient stimuli while they have found habitual responses in the medial aspects of PFC, amygdala and hippocampus (Phan et al., 2003). One possibility could be that the self-report is a habit-based responding of the individual to the drug cue and does not reflect the dynamic changes that are actually occurring in the brain and the brain might be processing drug cues to generate motivated behavior in a way that is not accessible to verbal report in the substance using individual. This may explain why individuals with SUD have a difficult time accurately predicting how they might respond in a “high risk” situation or how they might cope in those situations. Together, this lack of “insight” can contribute to the increased risk for relapse (H. Ekhtiari et al., 2016). This gap between subjective self-report of craving and brain response to the drug cues, due to the multi-dimensional nature and implicit aspects of craving, opens door for specific applications for the brain-based biomarkers of craving to provide additional values to the clinical practice.

One of the strengths of this study is the adoption of a discovery-replication model in testing hypotheses with brain imaging data. Serious concerns are mounting regarding the replicability of fMRI results (Poldrack et al., 2017). Using conventional exploratory methods in the whole brain or many ROIs can be associated with an increased risk of false positives that were not well controlled in many fMRI studies before (Eklund, Nichols, & Knutsson, 2016). In this study, we have used a discovery sample to find the areas that have significant effects in our analysis model, then used masks based on that analysis as a priori ROIs to test for replication in two independent samples. This kind of discovery-replication analysis pipeline will be helpful in reducing the risk for false positive results in the field. Replication of the same findings in other clinical populations and other labs across the world will be the next step for confirmation of our findings. To facilitate this confirmatory phase, we have shared our fMRI drug cue reactivity task (Hamed Ekhtiari et al., 2020) and our activation masks to help other labs to test the replication of the results in other research environments/groups (https://github.com/rkuplicki/LIBR_FDCR_Dynamic).

There are several limitations in this study. First, neither our study participants in the three sample populations nor our drug cues in the fMRI task have a clear label for being purely opioid or meth related. As it is obvious in the table 1 and also with anyone who has a clinical experience in addiction medicine, opioid and meth addiction have significant overlaps in both drug use rituals and also people who are affected in many parts of the world including US. Therefore, attributing the results of this study for being meth or opioid specific seems complex. However, as you can see in the table one, there are significant variations in terms of opioid and meth use profile among sample populations. Replication of response in this diverse set of clinical samples provide support for having a general response in both meth and opioid users instead of a drug specific behavior. Another limitation of this study is the limited temporal window of the experimental paradigm for 6.5 minutes and 4 blocks of cue exposure. This temporal window limits our sampling capacity to cover just rapid habituation response. Increase in the temporal window might bring potentials to detect areas with more subtle habituation response. Another important consideration in the interpretation of the results is the clinical context of the sample populations. All three sample populations are recruited from active treatment programs and participants were supposed to be back to the treatment setting right after the imaging session. As it could be expected in this setting, repeated exposure to the drug cues without any consequent drug use or expectation to drug use has not yielded in any sensitization response in subjective response or brain activations in our study. However, with change in the clinical context or drug use expectation, we can expect different outcomes.

This investigation is a one of the first studies on the temporal dynamics of responsiveness to drug cues. There are still many questions to be answered in future studies. One of the main remaining question is whether areas with or without habituation in response to drug cues are representing different cognitive constructs composing the craving phenomenon and how to modulate these cognitive processes with different cognitive or neuromodulatory interventions to stop the habituation process in the habituating areas or promote habituation in non-habituating areas. In the same context, we know that VMPFC, amygdala and VStr are highly connected areas (Paret et al., 2016), however the distinct role of these areas in the habituation response is still unclear. The clinical implication of these findings is another remaining question. Potentials for changing this dynamic response and their clinical significance and clinical value of this habituation as a potential biomarker or intervention development tool should be explored in the future.

In this preliminary study, we have explored the temporal dynamic response to the repeated presentation of drug cues among people with methamphetamine and/or opioid use disorders. We found that VMPFC, right amygdala and ventral striatum show a rapid habituation in response to drug cues, but not in the neutral cues. This habituation response was replicated in two other clinical populations. These areas will remain habituated to the drug cues in the second session of cue exposure few days later. This habituation in response could be a foundation to explore potentials for cue exposure interventions to modulate cue induced carving and reduce risks for potential relapse.

## Acknowledgement

Authors would like to thank Richard Turnham and Brad Collins in 12&12 addiction treatment center and Mimi Tarrasch in Women in Recovery for their invaluable support. Authors would also like to thank LIBR assessment team head Tim Collins, and his staff Brionne Sawyer, Katie Redman and Nigel Negm for their great work in the data collection for this project.

## Role of Authors

HE, RK and MPP developed the idea of the study, RK and HE have done the data analysis, HE has written the first draft of the manuscript and HE, RK and MPP have finalized and concluded about the submitted draft of the manuscript.

## Funding Agencies and Their Roles

Laureate Institute for Brain Research (LIBR), Warren K. Family Foundation, Oklahoma Center for Advancement of Science and Technologies (OCAST) grant #HR18-139 to HE and Brain and Behavior Foundation (NARSAD Young Investigator Award #27305 to HE)

