## Supplementary Material for "It is Never as Good the Second Time Around: Brain Areas Involved in Salience Processing Habituate During Repeated Drug Cue Exposure in Methamphetamine and Opioid Users"

### Supplements

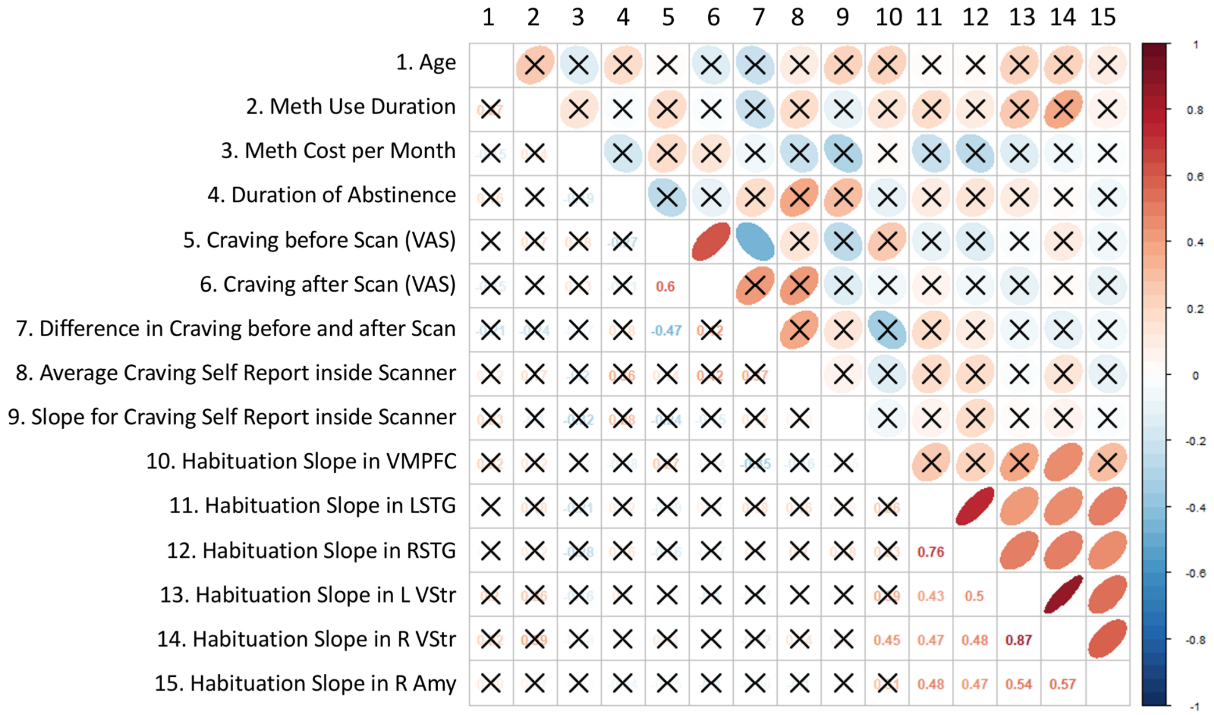


***Figure S1. Correlation matrix between clinical variables, craving self-reports and habituation slopes***. *Correlations that have not passed p value<0.05 threshold corrected for FDR are marked with cross. VAS: visual analogue scale, VMPFC: ventromedial prefrontal cortex, LSTG: left superior temporal gyrus, RSTG: right superior temporal gyrus, L VStr: left ventral striatum, R VStri: right ventral striatum, R Amy: right amygdala.*

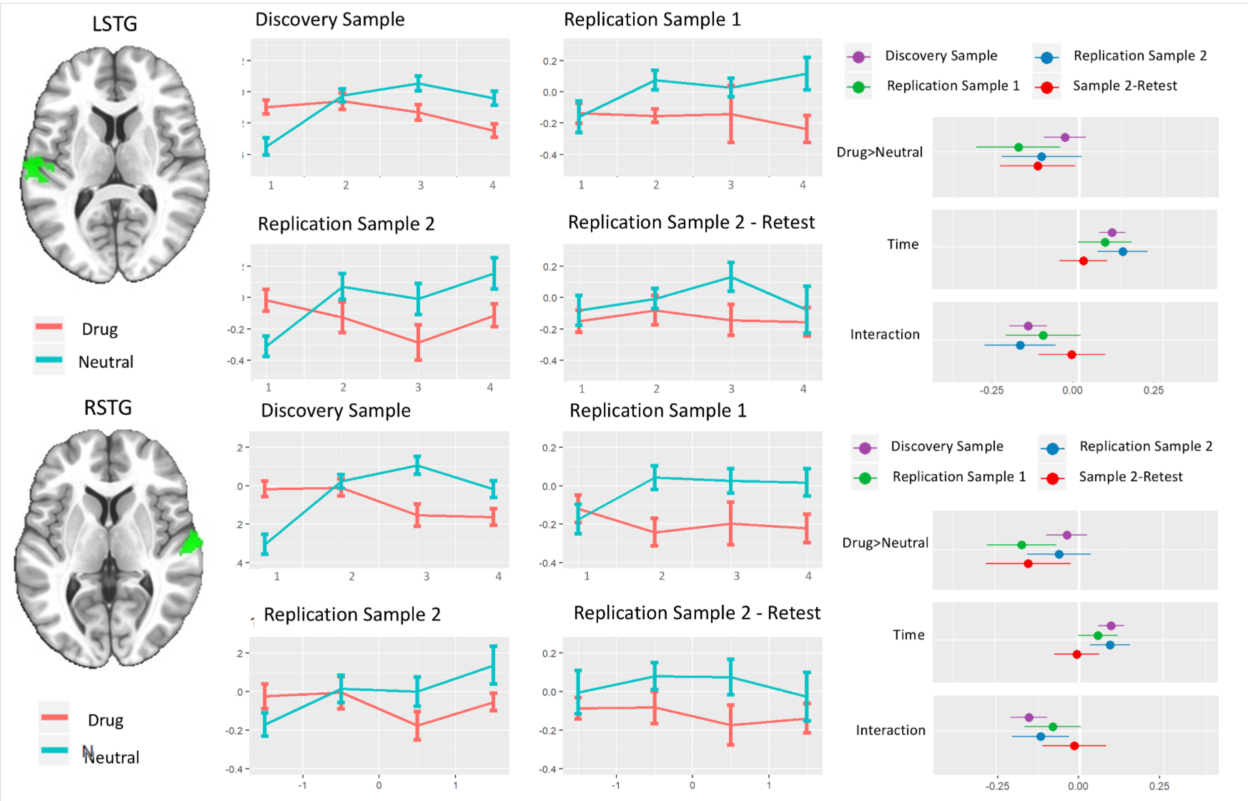


***Figure S2. Replication of Distinct Temporal Dynamic Response to Drug-related Cues Compared to Neutral Cues within Superior Temporal Gyrus (STG).*** *Middle panels show the temporal response to four drug and neutral blocks within the masks discovered from time by condition (drug/neutral) interaction in the discovery sample; error bars represent +- one standard error. The right panels show coefficients from four independent LME (linear mixed effect) models in three sample populations (one assessed twice) for the average beta estimates within the STG masks. Whiskers depict 95 percent confidence intervals.*
